# Laminin alpha 5 is Necessary for Mammary Epithelial Growth and Function by Maintaining Luminal Epithelial Cell Identity

**DOI:** 10.1101/2019.12.27.889451

**Authors:** Johanna I Englund, Hanne Cojoc, Leander Blaas, Alexandra Ritchie, Nalle Pentinmikko, Julia Döhla, Pauliina Munne, Manuel Patarroyo, Juha Klefström, Johanna Ivaska, Pekka Katajisto

**Affiliations:** Institute of Biotechnology, HiLIFE, University of Helsinki, Finland; Department of Bioscience and Nutrition, Karolinska Institutet, Stockholm, Sweden; Research Programs Unit/Translational Cancer Medicine, Medical Faculty and HiLIFE, University of Helsinki, Helsinki, Finland; Department of Microbiology, Tumor and Cell Biology, Karolinska Institutet, Stockholm, Sweden; Turku Bioscience Centre, University of Turku and Åbo Akademi University, Turku, Finland; Faculty of Biological and Environmental Sciences, University of Helsinki, Helsinki, Finland

**Keywords:** Laminin alpha 5, Extracellular matrix (ECM), Basement membrane (BM), Mammary gland, Luminal epithelial cell, Wnt4

## Abstract

Epithelial attachment to the basement membrane (BM) is essential for mammary gland development, yet the exact roles of specific BM components remain unclear. Here, we demonstrate that expression of distinct laminin α-isoforms by luminal and basal mammary epithelial cells enforces lineage identity that is necessary for normal mammary gland growth and function. Laminin α5 (LMα5) is mainly expressed by the luminal epithelial cells, and it is necessary for pubertal mammary gland growth, pregnancy induced gland remodeling, and for alveolar function. Adhesion to LMα5 containing laminin promotes luminal traits in both luminal and basal epithelial cells, and reduces progenitor activity of basal epithelial cells. Mechanistically, we show that *Lama5* loss interferes with differentiation of hormone receptor positive luminal cells, which results in reduced Wnt4 expression and defective crosstalk between luminal and basal epithelial cells during gland remodeling. Our results reveal a novel BM-mediated mechanism, which regulates mammary gland remodeling and function via specification of luminal epithelial cells.

## INTRODUCTION

The mammary gland is composed of a stroma-embedded epithelial network that undertakes the highly specialized function of milk secretion. Postnatal development of the mammary tissue occurs by branching and elongation of the epithelium into mammary stroma at the onset of puberty (Macias and Hinck 2012; Paine and Lewis 2017). Thereafter, the mammary gland is characterized by extensive proliferation, remodeling and apoptosis during the cycles of pregnancy and involution, which can occur multiple times throughout the reproductive lifespan. The fully formed adult mammary epithelium consists of a bilayered duct, where apico-basally polarized keratin 8/18-positive luminal epithelial cells face the ductal lumen and are surrounded by keratin 5/14-positive basal myoepithelial cells (Adriance et al. 2005; Macias and Hinck 2012). Luminal and basal cells exhibit distinct functions in the gland: specific luminal cells give rise to milk producing alveolar cells during pregnancy and lactation, while contractile basal cells aid in the milk flow (Adriance et al. 2005; Macias and Hinck 2012).

The mammary microenvironment contributes to various properties of mammary epithelial cells, including proliferation, survival, and differentiation (Inman et al. 2015). In particular, attachment to the specialized layer of the extracellular matrix (ECM) called the basement membrane (BM) is critical for cultured mammary epithelial cells and for mammary gland morphogenesis *in vivo* (Weaver et al. 1997; LaBarge et al. 2009; Inman et al. 2015). The BM acts as a physical barrier separating the epithelium from the stroma and as a scaffold supporting epithelial adhesion and tissue architecture (Yurchenco 2011). Moreover, BM regulates tissue homeostasis by supplying cells with growth factors and other signaling molecules and by regulating their availability to the cells (Yurchenco 2011).

Laminins (LM) are the main components of the BM that together with collagen IV form self-assembling networks, which serve as anchoring platforms for epithelial cells and provide various survival and differentiation signals (Hohenester and Yurchenco 2013). Laminins are heterotrimers consisting of *α*, *β*, and *γ* subunits, which are expressed in a tissue-specific and temporally controlled manner (Ahmed and Ffrench-Constant 2016). Several laminin isoforms have been detected in the mammary gland, and earlier studies suggest that Laminin-111 (LM-111; containing *α*1, *β*1, and *γ*1 subunits), LM-332 and LM-511/521 are the most common forms in the adult glands (Gudjonsson et al. 2002; Prince et al. 2002; Goddard et al. 2016), yet LM-211 and LM-411/421 are also found. A microarray profiling study showed that mRNA expression of *Lama1* and *Lama3* subunits is higher in a population of cells enriched for mammary stem cells (MaSC), whereas in mature luminal cells the expression of *Lama1* is reduced and *Lama5* is increased (Lim et al. 2010). In addition, recent data from single cell profiling studies confirm these results (Bach et al. 2017; Tabula Muris et al. 2018). However, what are the spatial and temporal expression patterns of these LMs within intact tissue and what are their exact roles in maintenance and function of luminal and basal mammary epithelial cells is unclear.

We set out to study the expression pattern of different LM isoforms in the mouse mammary gland during pubertal expansion, pregnancy and adult homeostasis *in vivo*. Different LM isoforms exhibited surprisingly discrete and cell type-restricted expression patterns in the mammary gland, and unexpectedly, we found strong expression of LMα5 in the luminal epithelial cells. We demonstrate that LMα5 produced by luminal cells enforces luminal identity, and is necessary for normal mammary gland growth and development both during puberty and pregnancy. Mechanistically, we show that LMα5 enhanced luminal differentiation is necessary for Wnt4 mediated interactions between luminal and basal cells during gland remodeling. Our results reveal that by orchestrating luminal cell specification during puberty and pregnancy LMα5 acts as a central microenvironmental effector of mammary gland function.

## RESULTS

### Laminins exhibit cell type specific expression patterns in the mammary epithelium

To explore the role of specific LMs in mammary epithelial cell (MEC) function, we first studied their expression during puberty, when the mammary epithelium expands and invades into the stroma via motile terminal end bud (TEB) structures (Macias and Hinck 2012). Using *in situ* hybridization (ISH) to detect the expression of *Lama1*, *3*, *4*, and *5* isoforms encoding the LM*α*1, *α*3, *α*4 and *α*5 subunits, we observed that *Lama1* and *3* were both expressed by the myoepithelial cells of established ducts (Fig S1) as well as in growing TEBs of pubertal virgin females (Fig 1a). In addition, *Lama3* was sporadically expressed in individual luminal epithelial cells of mature ducts (Fig S1, red arrow heads). In striking contrast, *Lama4* and *5* were strongly expressed by luminal epithelial cells, particularly in TEBs. *Lama4* expression by stromal cells was also evident, as previously reported (Petajaniemi et al. 2002) (Fig S1, asterisks). Next, we assessed the expression of *Lama1* and *Lama5* during pregnancy, where the epithelium undergoes substantial growth and differentiates to form alveolar structures. Similar to puberty, we observed a lineage-specific staining pattern with *Lama1* expression in basal and *Lama5* in luminal epithelial cells (Fig 1b). Finally, we combined ISH with keratin 8 (K8) or keratin 14 (K14) antibody staining to further asses the expression pattern of *Lama1* and *Lama5* (Fig 1c, Fig S1b). Confirming our findings, *Lama1* was mainly expressed by myoepithelial (K8-, K14+) cells, while *Lama5* was significantly enriched in luminal (K8+, K14-) epithelial cells (Fig 1d).

**Figure 1.**
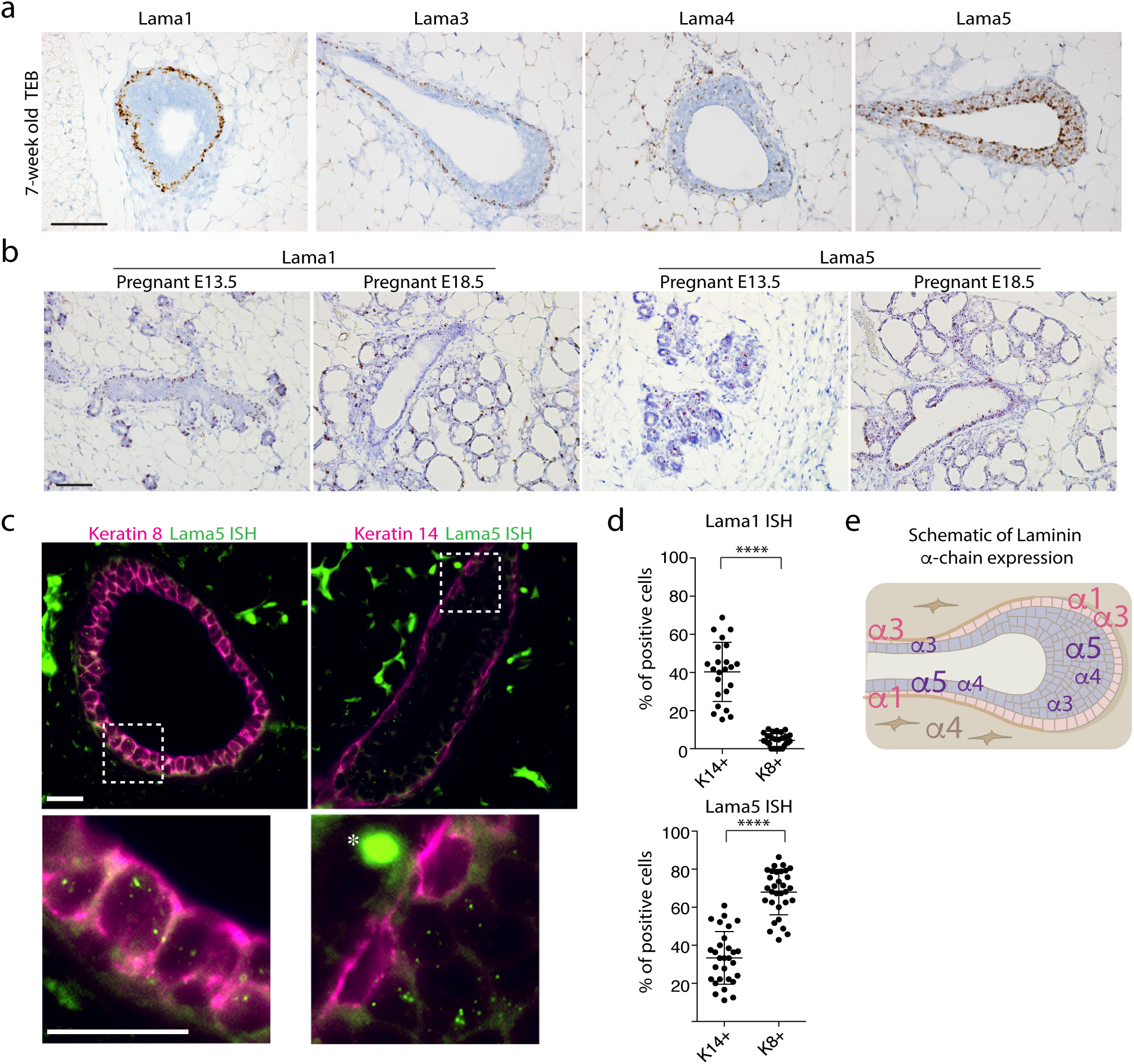
Expression of Laminin *α* chain isoforms in the mammary gland is cell type-specific. a) Representative images of RNA *in situ* hybridization performed with probes for *Lama1*, *3*, *4* and *5* on TEBs of 7-week old pubertal mammary glands. Scale bar 50 μm. b) Representative images of RNA *in situ* hybridization of mammary glands from E13.5 or E18.5 pregnant mice performed with probes for *Lama1*, and *5* on TEBs. Scale bar 50 μm. c) Representative images of *Lama5* RNA *in situ* hybridization (ISH) coupled with either K8 or K14 antibody staining. Scale bars 20 μm. d) Quantification of percentage of K14 or K8 positive cells showing either *Lama1* or *Lama5* expression. Data show mean ± SD. Two-tailed student’s t-test was used to compare groups. e) Schematic showing Laminin *α*-chain expression in the TEB and mature ducts of mouse mammary gland. Laminin *α*-chains expressed in the basal cell layer (pink) are depicted in light red and *α*-chains expressed in the luminal layer (light blue) are depicted in violet. Stromally expressed laminins are depicted in brown.

The surprisingly strong expression of specific laminins by the luminal cells prompted us to examine the localization of laminin proteins. Antibodies raised against LM*α*1, LM*α*4, and LM*α*5 resulted in unspecific staining, but a pan-laminin antibody revealed laminin localization mainly along the BM surrounding mature ducts and TEBs (Fig S1c), in line with previous reports (Paine and Lewis 2017). This suggests that the LM*α*5 expressed by the luminal cells is deposited into the BM and can affect BM composition. Indeed, using the *Lgr6-Cre-TdTomato* reporter mouse, where clones of exclusively luminal or basal cells can be visualized (Blaas et al. 2016), we detected contacts between luminal cells and the BM (Fig S1d). Such thin protrusions originating from luminal cells and reaching between basal cells to the BM have been demonstrated in earlier work (Smith and Medina 1988; Lafkas et al. 2013)

Taken together, these results indicate that LM*α* isoforms have distinct luminal, basal, and stromal expression patterns within the mammary gland (Fig 1e). Our findings also infer that both the luminal and basal epithelial cells could contribute to the laminin pool of the BM.

### Laminin *α*5 is required for pubertal mammary gland morphogenesis

Based on the surprising expression pattern of LM*α*5 in the mammary epithelium and the observed contacts with luminal cells and the BM, we set out to explore the physiological function of LM*α*5-containing laminins in mammary gland morphogenesis. To this end, we crossed mice harboring a conditional *Lama5* allele (Nguyen et al. 2005) with *K8-CreERT2* mice (Van Keymeulen et al. 2011) to allow tamoxifen-inducible deletion of *Lama5* in luminal epithelial cells. We determined the effects of *Lama5* deletion on pubertal mammary gland morphogenesis by injecting 3-week-old female *Lama5^fl/fl^;K8-CreERT2, Lama5^fl/+^;K8-CreERT2,* or *Lama5^+/+^;K8-CreERT2* mice with tamoxifen, and analyzed glands at 6 or 8 weeks of age (Fig 2a, Fig S2a). *Lama5* deletion in luminal MECs lead to delayed development of the mammary epithelium (Fig 2b), accompanied with diminished TEB structures (Fig 2c). In addition to the reduced TEB size, also the number of TEBs was reduced in mice with either heterozygous or homozygous *Lama5* deletion (Fig 2d). Correspondingly, total number of ductal ends and the extent of the epithelial network were also significantly reduced two weeks later at 8-weeks (Fig 2e-f, S2b). However, when *Lama5* was deleted in mature glands of 8-week old mice and analyzed at 12-weeks of age, no difference in the ductal end number was observed (Fig S2c). This data suggests that while *Lama5* is required for the mammary epithelium growth and morphogenesis during pubertal remodeling, loss of *Lama5* in adult mammary glands has little effect during homeostasis.

**Figure 2.**
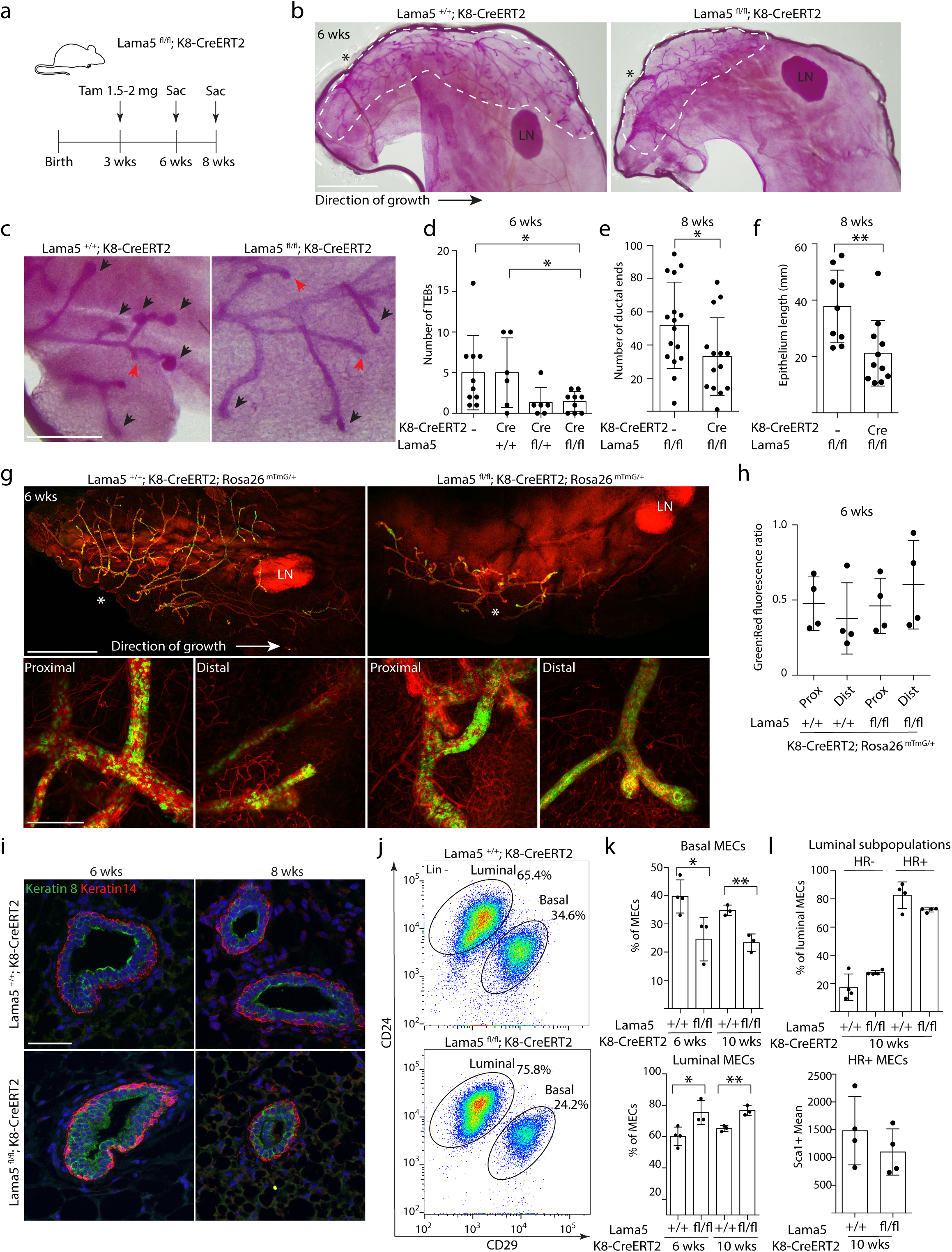
Luminal laminin *α*5 is required for pubertal growth of mammary epithelium. a) Schematic showing the outline of tamoxifen induction of K8-CreERT2 during puberty. b) Representative images of carmine alum-stained #4 mammary glands of 6-week old *Lama5^+/+^;K8-CreERT2* and *Lama^fl/fl^;K8-CreERT2* treated with tamoxifen. Line shows the outline of ductal growth and asterisk marks the beginning of epithelium. LN, lymph node. Scale bar 2 mm. c) Representative images of TEBs and ductal ends (marked by black arrow heads and red arrows, respectively) in 6-week old transgenic mice. Scale bar 0.5 mm. d) Quantification of the number of TEBs in 6-week old transgenic mice analyzed from #4 mammary glands. Data show mean ± SD. 1-2 glands were analyzed from individual animals. Two-tailed student’s t-test was used to compare indicated groups. e) Quantification of ductal ends and f) epithelium length in 8-week old transgenic mice analyzed from #4 mammary glands. Data show mean ± SD. Two-tailed student’s t-test was used to compare indicated groups. g) Representative images of #4 mammary glands of 6-week old *Lama5^+/+^;K8-CreERT2;R26^mTmG/+^* and *Lama5^fl/fx^;K8-CreERT2 R26^mTmG/+^* mice. Non-recombined cells exhibit red and recombined cells green fluorescence. Asterisk marks the beginning of epithelium. LN, lymph node. Scale bar 2 mm. Insets show close up images of proximal (closer to the beginning of epithelium) and distal parts of the duct, scale bar 200 µm. h) Quantification of green to red fluorescence ratio in proximal and distal parts of the duct in *Lama5^+/+^;K8-CreERT2;R26^mTmG/+^* and *Lama5^fl/fl^;K8-CreERT2 R26^mTmG/+^* mice. i) Representative immunofluorescence images of 6- and 8-week old *Lama5^+/+^;K8-CreERT2* and *Lama5^fl/fl^;K8-CreERT2* glands immunostained with K8 and K14 antibodies. Scale bar 50 µm. j) Representative CD24/CD29 FACS plot showing relative amounts of lineage marker negative luminal and basal MECs from 6-week old *Lama5^fl/fl^;K8-CreERT2* and control animals. Gating strategy for the same sample is shown in Fig. S2d. k) Quantification of the percentages of basal and luminal MECs from 6- and 10-week old *Lama5^fl/fl^;K8-CreERT2* and control mice. Data show mean ± SD from 3-4 individuals. l) Quantification of HR- and HR+ luminal subpopulations from 10-week old *Lama5^fl/fl^;K8-CreERT2* and control animals. Lower graph shows mean intensity of Sca1-positive luminal population from *Lama5^fl/fl^;K8-CreERT2* and control mice. Data show mean ± SD. Two-tailed student’s t-test was used to compare indicated groups.

To address whether gland development in *Lama5^fl/fl^;K8-CreERT2* mice was maintained by non-excised escaper clones of luminal cells, or whether *Lama5* deleted luminal cells contributed to the defective gland growth, we crossed *Lama5^fl/fl^;K8-CreERT2* mice with the *R26-mTmG* reporter line (Muzumdar et al. 2007). At six weeks of age and three weeks post tamoxifen injection, mammary glands in *Lama5*-targeted mice contained ubiquitous recombined cells in both proximal and distal ducts, indicating that *Lama5*-deficient cells did contribute to the blunted epithelial growth (Fig 2g-h).

To start addressing the cellular mechanisms blunting gland growth after *Lama5* loss, we first performed K8 and K14 antibody staining on *Lama5^fl/fl^;K8-CreERT2* and control glands at 6 and 8-weeks of age (Fig 2i). While K14-positive basal cells were unaltered, the K8 staining revealed aberrant organization and additional layers of luminal cells in the *Lama5*-targeted glands. As the histological analysis suggested that *Lama5*-lacking ducts may contain excess luminal cells, we next analyzed the frequency of CD29^hi^/CD24^+^ basal and CD29^low^/CD24^+^ luminal MECs in 6-week old *Lama5^fl/fl^;K8-CreERT2* and control glands with FACS. Three, and seven weeks after tamoxifen, glands with *Lama5* loss exhibited altered cellular ratios with a significant increase in luminal MECs (Fig 2j-k). Further analysis of the hormone receptor positive (HR+, Sca1^+^/CD49b^-^) and negative (HR-, Sca1^-^ /CD49b^+^) luminal subpopulations (Sleeman et al. 2007; Shehata et al. 2012) showed a modest decrease in HR+ cells that was interestingly accompanied with a reduction in the mean Sca1 expression in the remaining HR+ cells after *Lama5* loss (Fig 2l, S2d). To summarize, luminally produced *Lama5* was necessary for pubertal mammary gland growth, proper duct architecture, and for normal ratio of epithelial cells.

### Laminin *α*5 is needed for growth and differentiation of mammary epithelium during pregnancy and lactation

Next, we asked whether *Lama5* deletion affects the dramatic remodeling of the mammary glands during pregnancy, and the differentiation into milk producing alveoli. Growth and alveologenesis during pregnancy are driven by HR+ luminal progenitors, which in response to hormonal cues rapidly proliferate and generate luminal cells that form buds invading the ECM, and differentiate into the milk producing alveoli (Inman et al. 2015). We deleted *Lama5* in luminal cells of 8-week old female mice 1-2 weeks prior to pregnancy, and analyzed glands 17.5 days post coitum (dpc; Fig 3a). Alveolar development was dramatically reduced in *Lama5* deficient glands as evidenced by fewer and smaller alveoli compared to control glands (Fig 3a-b). Moreover, *Lama5*-deficient alveoli failed to form round and distinct lumen containing structures, and instead formed aggregates of luminal cells with marked reduction in surrounding Pan-laminin staining (Fig 3c). The disorganization of the epithelium was further illuminated by K8/K14 immunostaining demonstrating aberrant localization of both luminal and basal cells (Fig 3c).

**Figure 3.**
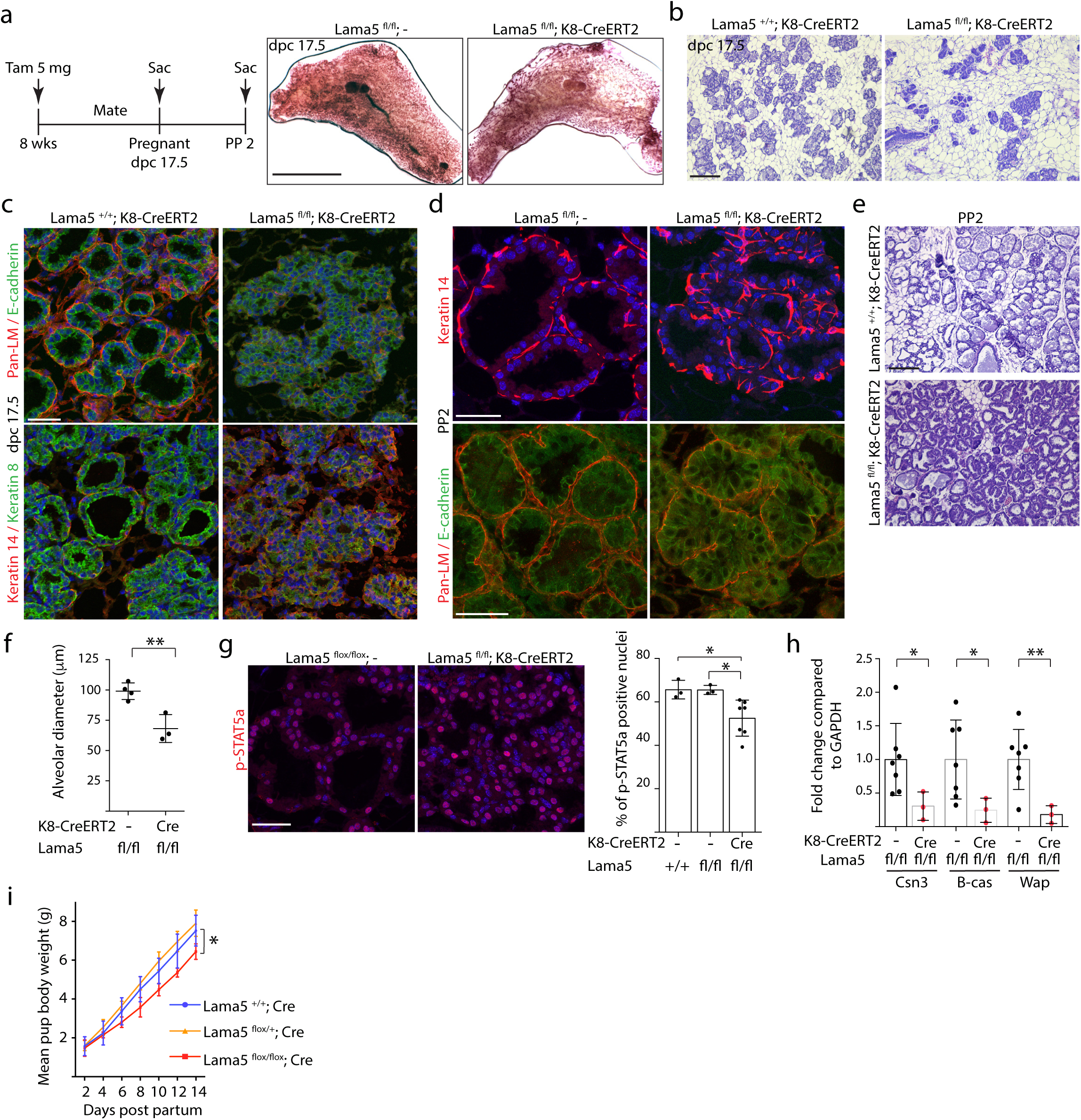
Laminin *α*5 is essential for pregnancy-induced growth and differentiation of mammary gland. a) Schematic showing the experiment outline of tamoxifen induction of K8-CreERT2 prior to pregnancy. Representative images of carmine alum-stained #4 mammary glands of 17.5 days post coitum (dpc) pregnant *Lama^fl/fl^;-* and *Lama^fl/fl^;K8-CreERT2* animals. Scale bar 10 mm. b) Representative HE images of dpc 17.5 pregnant *Lama5^+/+^;K8-CreERT2* and *Lama5^fl/fl^;K8-CreERT2* mice. Scale bar 200 μm. c) Representative immunofluorescence images of dpc 17.5 pregnant *Lama5^+/+^;K8-CreERT2* and *Lama5^fl/fl^;K8-CreERT2* glands immunostained with Pan-laminin and E-cadherin or K8 and K14 antibodies. Scale bar 50 μm. d) Representative immunofluorescence images of postpartum day 2 (PP2) mammary glands of *Lama^fl/fl^;-* and *Lama5^fl/fl^;K8-CreERT2* mice immunostained with K14 or Pan-laminin and E-cadherin antibodies. Scale bar 50 μm. e) HE staining of PP2 mammary glands of *Lama5^fl/fl^;-* and *Lama5^fl/fl^;K8-CreERT2* mice. f) Quantification of alveolar diameter of mice in e. Data show mean ± SD. Two-tailed student’s t-test was used to compare groups. g) Representative immunofluorescence images of p-STAT5a immunostaining in PP2 mammary glands of *Lama^fl/fl^;-* and *Lama5^fl/fl^;K8-CreERT2*, scale bar 50 μm, and quantification of the percentage of positive cells. Data show mean ± SD. h) qRT-PCR analysis of *Csn3*, *B-cas* and *Wap* expression compared to GAPDH in PP2 lactating *Lama^fl/fl^;-* and *Lama5^fl/fl^;K8-CreERT2* mice. Two-tailed student’s t-test was used to compare indicated groups. i) Mean body weight of pups during the first 14 days post partum nursed by *Lama5^+/+^, Lama5^fl/+^ or Lama5^fl/fl^;K8-CreERT2* mothers. Two-tailed student’s t-test was used to compare *Lama5^+/+^;K8-CreERT2* and *Lama5^fl/fl^;K8-CreERT2* groups at day 14.

We next examined how the aberrant alveologenesis caused by *Lama5* deficiency affects the capability of mammary epithelium to undergo functional differentiation during lactation, and analyzed mice exhibiting *Lama5* deletion at postpartum day 2 (PP2) (Fig 3a). Recapitulating the defects observed during pregnancy, *Lama5-* deficient alveoli were also significantly smaller during lactation, exhibiting disorganized tissue architecture with luminal cell aggregates and reduced Pan-laminin immunostaining (Fig 3d-f). Importantly, the *Lama5*-deficient glands exhibited fewer lipid droplets than control alveoli (Fig 3e), suggesting failed functionalization. To further examine whether *Lama5* is necessary for the functional differentiation of the epithelial cells, we stained glands for the phosphorylated form of STAT5a, which is increased in response to prolactin signaling in early lactation (Liu et al. 1996). Phospho-STAT5a staining was significantly decreased in Lama5 deficient glands at PP2 (Fig 3g), and accordingly, expression of milk protein genes *Csn3*, *B-cas* and *Wap* was also significantly reduced (Fig 3h). Finally, as a functional readout of milk production we measured the weight of pups of mothers lacking either one or two alleles of *Lama5* over 14-day time period starting at day 2 and compared to pups nursed by control mothers. Demonstrating the defective mammary gland function, pups nursed by mothers with biallelic luminal *Lama5* deletion gained weight significantly slower than other groups (Fig 3i).

Taken together, these data demonstrate that *Lama5* is required for proper tissue architecture and functional differentiation of mammary glands during pregnancy and lactation. Altogether our findings with the *Lama5* mouse model demonstrate an important *in vivo* role for *Lama5* in luminal MECs during key stages of mammary gland development and function.

### Adherence to Laminin *α*5 enhances luminal gene expression, and suppresses basal gene expression and function

Our results demonstrate that luminal LM*α*5 is critical for the growth and functional differentiation of the mammary epithelium and previous results have shown that adhesion to LM-111 is critical specifically for basal epithelial function (Gudjonsson et al. 2002). Thus, we next wanted to address how adhesion to *Lama1-* and *Lama5*-containing laminins in the mammary epithelial microenvironment affects luminal and basal epithelial cell properties at the molecular level. To mimic the effects of major laminin components of the mammary BM, we coated culture plates with recombinantly produced laminins LM-111 and LM-521 containing the laminin *α*1 and *α*5 subunits respectively, and cultured freshly isolated CD29^low^/CD24^+^ luminal or CD29^hi^/CD24^+^ basal mouse mammary epithelial cells on these laminins (MMECs; Fig S3a-b, Fig 4a). We observed that luminal and basal MMECs grown on LM-111 had typical morphology - cuboidal for luminal, and more elongated for basal epithelial cells (Fig 4b). Interestingly, both cell types exhibited flat and spread morphology on LM-521 (Fig 4b) suggesting that the LM-521 provides a particularly adhesive substratum for mammary epithelium. We confirmed these results with a human basal mammary epithelial cell line (HMEC-FL2), and observed that cells grown on LM-111 had a typical basal epithelial morphology whereas cells on LM-521 were extensively flat (Fig 4c). Scanning electron microscopy of the HMEC-FL2s on LM-111 and LM-521 adhered cultures corroborated these findings (Fig 4c).

**Figure 4.**
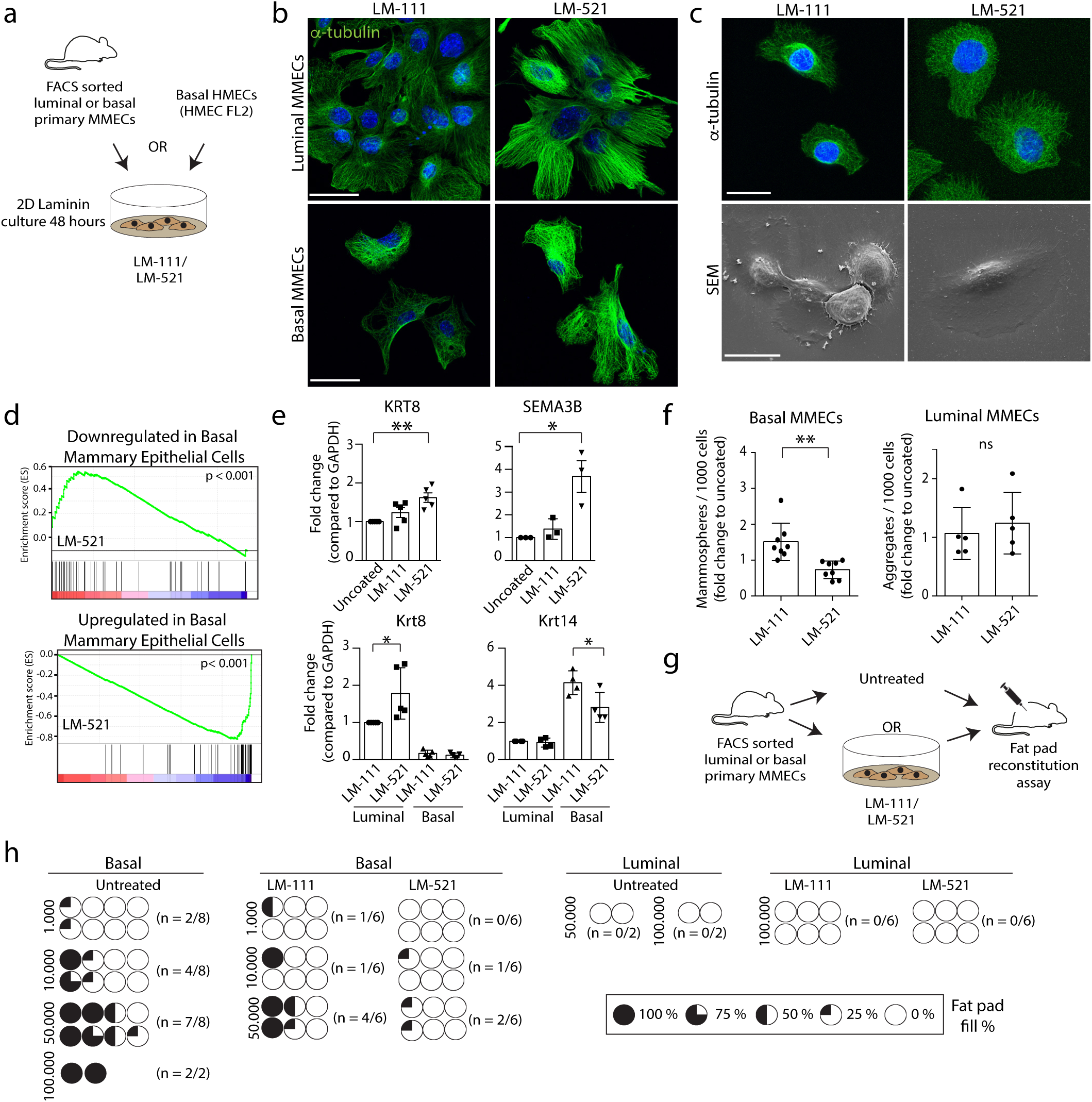
Adhesion to specific Laminins affects luminal and suppresses basal gene expression and basal progenitor activity. a) Schematic showing the outline of the laminin preadhesion assay, in which primary MMECs or HMECs are cultured on 2D LM-111 or LM-521 matrix for 48h. b) Representative immunofluorescence images of FACS sorted luminal and basal MMECs grown on indicated laminins and immunostained with *α*-tubulin. Scale bar 50 μm. c) Immunofluorescence images of *α*-tubulin and scanning electron (SEM) micrographs of HMECs grown on the indicated laminins. Scale bars 20 μm. d) GSEA plots showing correlation between the signature of HMECs grown on LM-521 and previously reported gene sets either downregulated in basal MECs (upper panel) or upregulated in basal MECs (lower panel). e) qRT-PCR analysis of KRT8 and SEMA3B expression compared to GAPDH in HMECs grown on indicated laminins for 48h and qRT-PCR analysis of Krt8 and Krt14 expression compared to GAPDH in basal (CD29^hi^/CD24^+^) and luminal (CD29^low^/CD24^+^) MECs pre-adhered on LM-111 or LM-521 for 48h. Data show mean ± SD. Two-tailed student’s t-test was used to compare indicated groups. f) Mammosphere or aggregate forming frequency of FACS sorted luminal and basal primary MMECs pre-adhered on the indicated laminins 14 days earlier. Data show mean ± SD from independent experiments. g) Schematic showing laminin pre-adhesion assay outline, in which FACS sorted primary MMECs are used for fat pad reconstitution assay either without treatment or after 48 hour of culturing on LM-111 or LM-521 matrix. h) Limiting dilution fat pad reconstitution of basal (CD29^hi^/CD24^+^) and luminal (CD29^low^/CD24^+^) MECs pre-adhered on LM-111 or LM-521 or untreated controls. Filling of the circles represents the extent of reconstitution (0, 25, 50, 75 or 100%) and each circle represents one gland.

Next, we performed RNA sequencing of the basal HMEC FL2 cells grown on either LM-111, -521 or on uncoated control plates for 48 hours. Culture on LM-111, which is expressed by basal cells, induced expression changes only in 8 genes, when compared to cells from uncoated plates (Fig S3c). In contrast, LM-521 adhesion induced expression changes in 944 genes. Hallmark gene sets related to metabolism and MTORC1 signaling were enriched on LM-521 (Fig S3d, p<0.05) and gene sets related to e.g. Myc targets and the G2M checkpoint were downregulated, as shown by Gene Set Enrichment analysis (GSEA) (Subramanian et al. 2005). This was surprising, as we did not observe significant changes in cell proliferation on any of the laminins (Fig S3e-f). Interestingly, however, the gene sets most significantly enriched in LM-521 grown cells also contained the genes typically expressed in luminal cells (Huper and Marks 2007), whereas genes typically expressed by basal cells (Huper and Marks 2007) were down regulated (Fig 4d). The increased expression of luminal markers in LM-521 grown basal HMECs was validated with qRT-PCR (Fig 4e). Intriguingly, adhesion to LM-521 enhanced K8 expression also in FACS-sorted luminal MMECs, and while LM-521 was not sufficient to induce K8 expression in primary basal MMECs, it resulted in significant decrease in basal marker Keratin 14 (K14) (Fig 4e). Taken together, these results show that adhesion to LM-521 enforces luminal gene expression in MECs along with a decrease in expression on basal markers.

Our results implied that adhesion to different laminins can drive a conversion of basal cell identity to a luminal direction and thus we wanted to further test whether adhesion to luminal Laminin α5 regulates also basal cell function. We first tested how attachment to various laminins affects mammosphere forming capacity of luminal and basal MECs after 48-hour preadhesion to either LM-111 or LM-521. Mammosphere assay allows quantitative assessment of progenitor cell clonogenicity of the mammary epithelium (Shaw et al. 2012) *in vitro*. As expected, preadhesion of basal MECs to LM-111 increased mammosphere formation, but in striking contrast, preadhesion to LM-521 decreased mammosphere formation significantly (Fig 4f). As a control, luminal cells formed non-spheroidal aggregates at a constant frequency irrespective of preadhesion (Fig 4f). Similar results were obtained with unsorted MMECs and basal HMECs, where LM-111 resulted a minor increase, while LM-521 significantly decreased mammosphere formation (Fig S3g-h). Finally, we performed a limiting-dilution mammary reconstitution assay to explore the *in vivo* effects of laminin pretreatments on progenitor activity of basal cells (Shackleton et al. 2006; Stingl et al. 2006; Sleeman et al. 2007). Sorted basal and luminal cells were both preadhered on either LM-111 or LM-521 for 48 hours and 1000, 10000, or 50000 cells were transplanted per gland (Fig 4g). Up to 100000 freshly isolated basal and luminal cells were directly transplanted as controls. As expected, transplanting untreated basal cells resulted in outgrowths with increasing frequency, while luminal cells yielded no outgrowths (Fig 4h). In agreement with our previous results, LM-111 preadhered basal cells formed outgrowths more often than LM-521 preadhered basal cells (Fig 4h), and 50000 LM521 preadhered cells were required to achieve an outgrowth efficiency comparable to 1000 LM-111 preadhered cells. As expected, LM-111 or LM-521 preadhered luminal cells did not form outgrowths. (Fig 4h). Consequently, adhesion to specific laminin isoforms critically modulates gene expression and function including tissue reconstitution capacity of basal MECs.

### *Lama5*-deficient luminal MECs are unable to support basal cells

Our data suggested that Laminin *α*5 adhesion is a strong inductive signal for luminal lineage in mammary epithelial cells. Ductal elongation and alveologenesis requires interplay between the luminal and basal cells, and we hypothesized that such interactions may be defective in Lama5 lacking epithelium where luminal differentiation is compromised. As an example of luminal-basal interactions, progesterone-induced Wnt4 secretion by the HR+ luminal epithelial cells together with the R-spondin 1 produced by the HR-luminal cells is necessary for maintenance of basal epithelial cell function and hence can affect the growth of mammary epithelium as a whole (Brisken et al. 2000; Cai et al. 2014; Rajaram et al. 2015). Intriguingly, our data showing reduced Sca1 expression in HR+ luminal cells after *Lama5* deletion, and recent data demonstrating that *Lama5* is made mostly by the HR+ cells (Tabula Muris et al. 2018), jointly suggest that HR+ cells could be defective in the Lama5 lacking glands. Thus, we investigated the expression of Wnt4 in FACS sorted luminal and basal MECs from 8-week old *Lama5^fl/fl^* and control mice and observed that Wnt4 expression was indeed significantly decreased in *Lama5* deficient luminal epithelial cells (Fig 5a).

**Figure 5.**
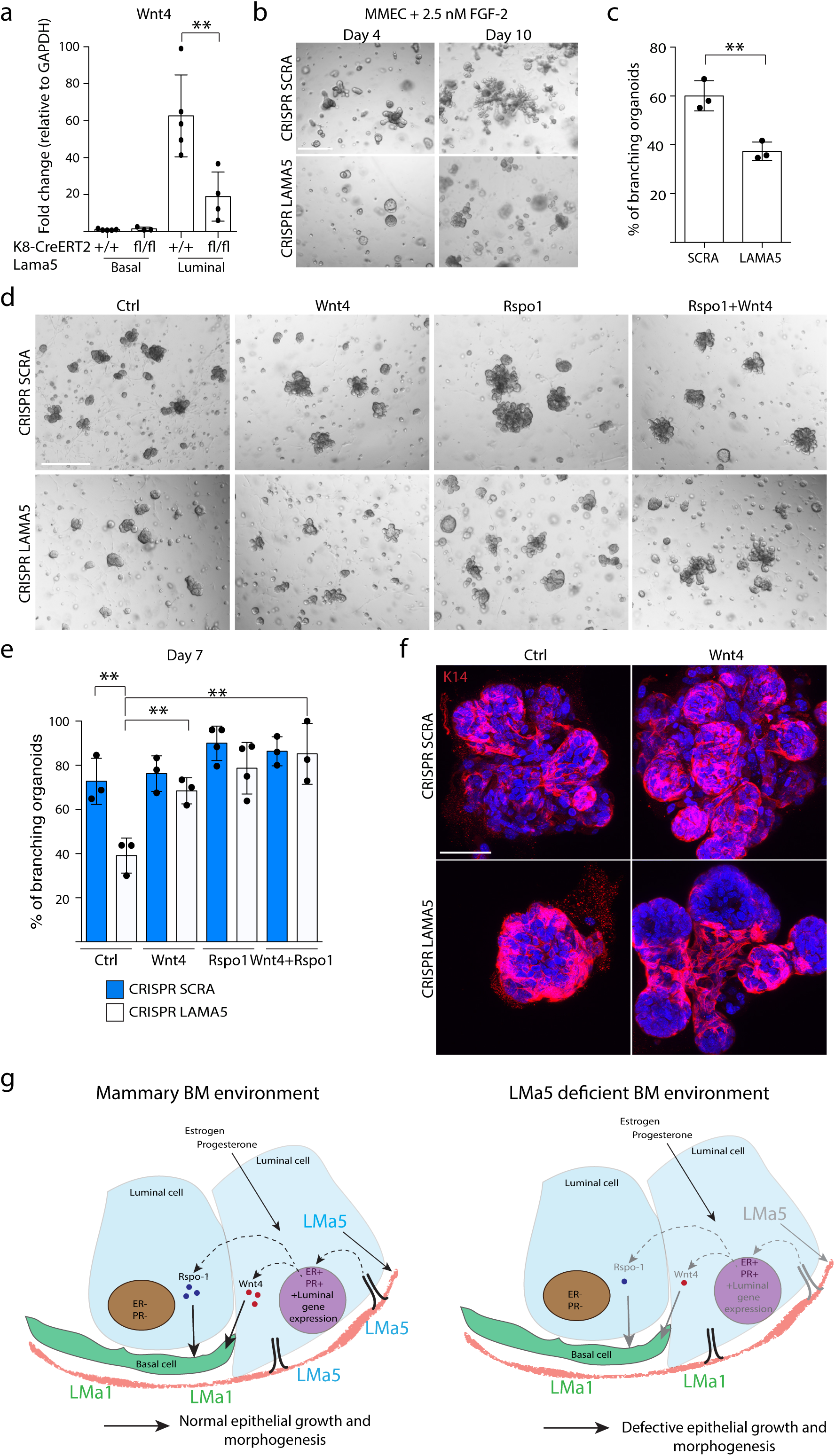
Luminal MECs with *Lama5* deletion are unable to support basal MECs due to defective Wnt signaling. a) qRT-PCR analysis of Wnt4 expression compared to GAPDH in basal (CD29^hi^/CD24^+^) and luminal (CD29^low^/CD24^+^) MECs FACS sorted from 8-10-week old *Lama5^+/+^;K8-CreERT2* and *Lama5^fl/fl^;K8-CreERT2* glands treated with tamoxifen starting from 3 weeks of age. b) Primary MMECs carrying either LAMA5 or SCRA CRISPR guides grown in Matrigel in the presence of 2.5 nM FGF-2 for 4 or 10 days. Scale bar 100 μm. c) Quantification of the percentage of branching organoids at day 10 grown as in b. Data show mean ± SD of 3 independent experiments. Two-tailed student’s t-test was used to compare groups. d) Primary MMECs carrying either LAMA5 or SCRA CRISPR guides grown in Matrigel in the presence of 2.5 nM FGF-2 and either Wnt4, Rspo1 or both ligands for 7 days. Scale bar 100 μm. e) Quantification of the percentage of branching organoids at day 7 grown as in d. Data show mean ± SD of 3 independent experiments. Two-tailed student’s t-test was used to compare indicated groups. f) Representative K14 immunofluorescence staining of primary MMECs organoids carrying either LAMA5 or SCRA CRISPR guides and treated as in d and grown for 7 days. Scalebar 50 μm. g) Schematic showing how production of LM*α*5 by luminal MECs supports the luminal cells and enhances luminal gene expression including Wnt4 for normal epithelial growth and morphogenesis. Deficiency of LM*α*5 in the BM environment lacking in turn leads to reduced luminal gene expression and defective epithelial growth.

Next, we tested whether addition of exogenous Wnt ligands can rescue the *Lama5-* dependent growth defect. Ductal growth by the TEB structures can be modeled *in vitro* with FGF-2 supplemented 3D cultures, where MEC organoids form branching structures (Ewald et al. 2008). First, we examined whether LM*α*5 is required for duct growth elongation of epithelial structures in this setting, and infected wild-type primary MMECs with lentiviruses carrying CRISPR/Cas9 and guide RNAs against *Lama5* or a non-targeting control sequence (Fig S4a). *Lama5* editing resulted in significant reduction in organoid branch formation and elongation (Fig 5b-c) similar to luminal *Lama5* deficiency in vivo. Subsequently, we added either Wnt4 or R-Spondin 1 separately, or in combination, into the FGF-2 induced 3D cultures. We observed that addition of either of these ligands increased branches in wild-type MMEC 3D organoids (Fig 5d), yet no increase in the overall branching frequency was observed (Fig 5e). However, in *Lama5*-deficient organoids there was a clear increase in branching of organoids with Wnt4 treatment, while combination of Wnt4 and R-Spondin 1 resulted in a largest increase in organoid branching compared to either ligand alone (Fig 5e). To test whether activation of Wnt signaling in general can phenocopy the observed effect, we tested whether Wnt3a and Wnt5a can also enhance branching morphogenesis in *Lama5*-deficient organoids. Previous results have also suggested that although Wnt3a is not produced in the mammary epithelium, it can augment basal cell function (Zeng and Nusse 2010), and Wnt5a can increase basal mammary spheroid growth (Kessenbrock et al. 2017). In accordance, addition of either of the ligands alone or in combination with R-Spondin 1 resulted an increase in branching in the *Lama5*-deficient organoids (Fig S4b). Conversely, inhibition of Wnt production by porcupine inhibitor IWP-2 decreased the amounts of branching organoids, yet it did not completely inhibit branch formation (Fig S4b).

Finally, to evaluate whether exogenous Wnt ligands restore the cell-type interaction dependent branching morphogenesis program in *Lama5*-deficient organoids, instead of e.g. activating basal cell proliferation, we immunostained control and *Lama5*-deficient, Wnt-treated organoids with K14 (Fig 5f). We observed, that even though *Lama5*-lacking organoids exhibited both K14+ and K14-cells, cell types were poorly organized and formed no branches. Strikingly, upon Wnt4 treatment the *Lama5*-lacking organoids formed elongated branches that were covered by K14-positive basal cells similarly to control organoids (Fig 5f). Furthermore, in line with our findings *in vivo* (Fig 2,3), *Lama5*-deficient organoids were not defective in proliferation, but Wnt4 normalized the pattern of proliferation in *Lama5*-lacking organoids as observed with immunostaining for Ki-67 (Fig S4e). These data indicate that Wnt4 reinstates the morphogenic branching program by facilitating the interplay between basal and luminal epithelial cells in the *Lama5*-lacking epithelium.

In conclusion, we show that adhesion to LM*α*5 produced by the luminal epithelial cells is instructive for luminal lineage gene expression and necessary for differentiation of HR+ cells, which can guide the remodeling of the whole mammary epithelium via Wnt4 secretion (Fig 5g).

## DISCUSSION

The mammary ECM is essential for normal physiology and pathology of the mammary gland (LaBarge et al. 2009; Maller et al. 2010; Goddard et al. 2016). However, the exact functions of specific BM constituents in mammary gland physiology have been unclear. We investigated the expression patterns of different laminins and explored how laminin adhesion affects luminal or basal MEC function *in vitro* and *in vivo.* Here, we show that BM laminin subunits *α*1, *α*3, *α*4 and *α*5 exhibit distinct gene expression patterns in the luminal, basal, and stromal compartments of the mammary gland. We discovered that *Lama5*, which is one of the most common laminin isoforms expressed in various mouse tissues (Yang et al. 2009), is predominantly expressed in the luminal MECs in mature glands and by centrally located cells within TEBs of the growing epithelium.

By deleting of LM*α*5 in the luminal cells at various stages, we found that it has a critical role during pubertal growth and pregnancy induced remodeling and functionalization. *Lama5* deficiency affected the ductal epithelial organization resulting in unorganized luminal cell aggregates within mammary ducts in both puberty and pregnancy. Our results imply that the *Lama5^-/-^* luminal cells proliferate but fail to maintain contacts with the BM, and consequently may result in the observed accumulation of unorganized cell aggregates filling the gland lumen. The defective organization may also reflect the loss of apico-basal polarity of luminal epithelial cells, previously shown to underlie defects in mammary morphogenesis (McCaffrey and Macara 2009; Akhtar and Streuli 2013). This can be further attributed to the loss of luminal epithelial cell contact with the BM, as luminal-ECM interaction is necessary for alveolar differentiation and, thus, functionalization and milk secretion (Roskelley et al. 1994; Streuli et al. 1995; Inman et al. 2015). Indeed, mammary epithelium lacking the adhesion receptor *β*1-integrin exhibits defective apico-basal polarization, surplus luminal cells within the ducts and impaired alveolar cell function (Naylor et al. 2005; Akhtar and Streuli 2013). Another possible cause for accumulation of extra luminal cells is defective luminal differentiation from progenitors to mature ductal and alveolar cells. In mammary epithelium with luminal-specific *Gata3* transcription factor deletion, there is a striking defect in epithelial growth and differentiation during puberty and pregnancy due to an increase in the pool of luminal progenitor cells (Asselin-Labat et al. 2007). Similarly, loss of another luminal transcription factor *Elf5* results in an alveolar differentiation defect and accumulation of luminal progenitors In accordance, we observed a decline in the mean intensity of Sca1 and Wnt4 in the *Lama5* lacking epithelium. Sca1 expression is used to identify ER*α* positive mature luminal cells and progenitors (Sleeman et al. 2007; Shehata et al. 2012) and Wnt4 is produced by hormone receptor positive cells (Brisken et al. 2000; Cai et al. 2014; Rajaram et al. 2015). As we detected only minor changes in the proportions of HR- and HR+ luminal cell pools, our data implies qualitative changes in the *Lama5* deficient HR+ luminal compartment. These data also suggest an interesting hypothesis that cell polarity and luminal differentiation are coupled downstream of BM adhesion, yet further experiments are needed to address this question.

During puberty and pregnancy, the growth of the epithelium occurs in a coordinated manner requiring interplay between both luminal and basal cell types (Macias and Hinck 2012; Inman et al. 2015). We sought to determine whether *Lama5* deficiency in luminal cells impairs mammary growth and remodeling cell-autonomously or via basal cell interactions. Previous studies have established Wnt signaling as an important mediator of the interplay between luminal and basal epithelial cells (Zeng and Nusse 2010). Wnt4 and R-spondin1, which are respectively produced by the HR+ and HR-luminal epithelial cells, act synergistically on basal cells during remodeling (Cai et al. 2014; Rajaram et al. 2015). We show that *Lama5* lacking luminal epithelial cells produce less Wnt4, and exhibit reduced branch formation in an *in vitro* 3D cell culture assay. This defect can be rescued by addition of exogenous Wnt4 and R-spondin1, and conversely, branch formation was inhibited similarly to *Lama5* deficiency when Wnt ligand secretion was inhibited in wild-type cells. While altered luminal-basal interactions at least partly explain the defective remodeling of *Lama5*-deficient glands, how LM*α*5 adhesion regulates Wnt4 signaling remains an open question. One possibility is that LM*α*5 adhesion leads to increased Wnt4 expression in luminal epithelial cells via enforcing luminal differentiation, more specifically of the ER+ lineage. Wnt4 has been shown to be regulated by progesterone (Brisken et al. 2000), which in turn is a target for ER*α* signaling (Macias and Hinck 2012) that could be orchestrated by LM*α*5 adhesion. In conclusion, our data reveals a mechanism through which BM adhesion regulates mammary epithelial growth as a whole via induction of Wnt4.

Finally, our data suggests a concept, in which lineage specification of mammary epithelium is controlled by autocrine production of a BM laminin microenvironment. Our results and previous studies demonstrate that LM*α*1 is the main laminin subunit produced by basal epithelial cells (Gudjonsson et al. 2002) and adhesion to LM*α*1 is essential for basal epithelial cell function. Interestingly, our results show in a complementary fashion a novel role for LM*α*5 adhesion, which exerts a strong luminalizing signal to mammary epithelial cells. Adhesion to LM*α*5 containing laminins enforced luminal gene expression even in basal epithelial cells, and repressed their *in vivo* mammary reconstitution capability. These data also suggest that similarly to skin (Morgner et al. 2015), the various laminin isoforms in the ECM of mammary epithelium need to be carefully balanced in order to optimally support both basal and luminal lineages. Additionally, our data suggest that LM*α*5 could be a novel upstream regulator of luminal epithelial cell differentiation. In the light of lineage tracing studies demonstrating that segregation of the mammary epithelial lineages occurs at a specific time during embryonic development (Lilja et al. 2018; Wuidart et al. 2018), it will be interesting to probe the role of laminin microenvironment in this process, and whether laminins aid the separation of luminal and basal epithelial cell identities in the developing gland.

## MATERIALS AND METHODS

### Animals

Animal studies were approved by the National Animal Ethics Committee of Finland (ELLA) and conducted in the Laboratory animal center (LAC) of the University of Helsinki under institutional guidelines or approved by the ethics committee of the Board of Agriculture, Experimental Animal Authority, Stockholm, Sweden. Female wild type C57BL6/RccHsd donor and recipient mice for fat pad reconstitutions were obtained from Envigo. The *Lgr6*-*EGFP*-*IRES-Cre^ERT2^ Rosa26-LSL-tdTomato*, *Lama5 fl*, *K8-CreERT2 and R26-mTmG* mouse lines have been previously described (Nguyen et al. 2005; Muzumdar et al. 2007; Van Keymeulen et al. 2011; Blaas et al. 2016). Genotyping of the animals was performed using previously described primers. Excised *Lama5* allele was detected with following primers Fwd: 5’-ACCTGGCTTTGACGGTCCT-3’ Rev: 5’-GTTGAAGCCAAAGCGTACAGCG-3’. For Lgr6-Cre induction, a single dose of 1 mg tamoxifen (Sigma) in 50 µl sunflower oil (Sigma) was injected intraperitoneally in 2-week-old mice. For K8-CreERT2 induction, a single dose of 1.5-2 mg tamoxifen in corn oil (Sigma) was injected intraperitoneally in 3-week-old mice, and 5 mg of tamoxifen in 8-week-old mice.

### RNA in situ hybridization

Gene expression analysis by RNA *In Situ* hybridization was performed using the RNAscope 2.5 HD Reagent Kit-Brown or RNAscope Multiplex Kit (Advanced Cell Diagnostics Srl) according to manufacturer’s instructions. 5 μm thick paraffin embedded sections were deparaffinized, rehydrated and pretreated with RNAscope Target Retrieval Reagents. A barrier was created around each tissue section on the slides, dried overnight, and used the following day. Pretreated samples were hybridized with either LM *α*1, *α* 3, *α*4 or *α*5 probes (custom made by Advanced Cell Diagnostics), or with positive or negative control probes (Ubiquitin or DapB, respectively) for 2 hours at 40 °C. Thereafter, signal amplification hybridization was performed, followed by detection with DAB and counterstain with hematoxylin. Alternatively, when Multiplex Fluorescent assay was used, after signal amplification hybridization the slides were immunostained with primary antibodies (K8 1:1000, TROMA-1 Developmental Studies Hybridoma Bank, K14 1:2000, Covance) for 1 hour. Thereafter samples were washed 3 times with PBS, 5 minutes each, and incubated with secondary antibodies in 10% NGS in IF Buffer for 1-2 hours. Slides were next washed 3 times followed by counterstaining of nuclei with 4’6-diamidino-2-phenyindole (DAPI) and mounted. Samples were imaged with Zeiss Axio Imager.M2 and AxioCam HRc camera using Plan Neofluar 20X (NA 0.5) or Plan Apochrom 63X (NA 1.4, oil) objectives and Zen software or with Nikon Inverted widefield system with 60x Plan apochromat objective (NA 1.2, water) and Andor Xyla 4.2+ camera and NIS elements software.

### Wholemount stainings

Mammary gland tissue samples for wholemount staining were fixed in 4 % PFA overnight. For wholemount staining, #4 inguinal mammary glands were stained for several hours in carmine-alumn staining solution (2% w/v Carmine, Sigma; 5 % w/v Aluminium Potassium Sulfate, Sigma). After the desired color had developed, glands were mounted to glass coverslips and imaged.

For wholemount imaging and immunostaining of *Lgr6-CreERT2*;*Rosa26-tdTomato* or *Lama5 fl;K8-CreERT2;Rosa26-mTmG* mammary glands the dissected mammary glands were pre-fixed in 4% PFA for 30–60 minutes at RT to preserve native fluorescence. For wholemount immunostaining, the tissue was permeabilized with 2 washes of PBS0.5T (PBS + 0,5% v/v Triton X-100) for 1h and the samples were incubated overnight at 4°C with primary antibody against Keratin 14 (Abcam, 1:500) diluted in PBD0.2T (PBS + 1%BSA + 1% v/v DMSO + 0.2% v/v Triton X-100). The next day, they were washed 4 times with PBD0.2T for 1h. Tissues were incubated with secondary antibodies plus DAPI (4’,6-diamidino-2-phenylindole; Roche) at 4°C overnight. After 4 washes with PBD0.2T for 1h, tissues were optically cleared in 80% glycerol in PBS at room temperature. The stained mammary glands were mounted between 2 cover slips and sealed with silicone. Imaging of the wholemounts was using a Zeiss LSM710 confocal microscope. A dry lens (5x, EC Plan Neofluar, NA 0.16) or water immersion lense (20X W Plan Apochromat NA 1.0 or 40x C-Apochromat, NA 1.2) was used to record optical sections at 512 x 512, 1024 x 1024 or 2048 x 2048 pixels and data were processed using Zeiss’ ZEN software. Images were converted to RGB images with ImageJ software.

### Morphometric analysis of the mammary gland

For quantitation of the number of terminal end buds (TEB) and ductal ends in the mammary glands and the length of the epithelium, #4 mammary glands stained with carmine-alumn wholemount were imaged with Leica S9i stereomicroscope equipped with integrated 10MP camera. The amount of TEBs and ductal ends was quantitated from the images based on morphology. Length of the epithelium was measured from the same images in ImageJ software by using the line tool. The length of the whole epithelial network was recorded.

### Quantification of fluorescence in mammary gland wholemounts

Mammary gland wholemounts of *Lama5^+/+^;K8-CreERT2;Rosa26-mTmG* and *Lama5^fl/fl^;K8-CreERT2;Rosa26-mTmG* were imaged for 3-5 proximal and distal areas of the mammary epithelium. Z-stacks were converted to Z projection using sum slice method. Regions of interest (ROI) were drawn to encompass the epithelium and three spots to quantify the background fluorescence. The integrated density of green and red channels of ROIs was measured, and mean background fluorescence was subtracted from the epithelial ROIs. From these values the green to red fluorescent ratio was calculated for both proximal and distal areas. Data is shown as mean per mammary gland.

### Tissue Immunostainings

Mammary gland tissue samples for immunostaining were fixed in 4 % PFA overnight and embedded in paraffin for sections. For hematoxylin and eosin (H&E) stainings, 5 μm thick sections were used. Stainings were performed in the Finnish Center for Laboratory Animal Pathology (FCLAP) at University of Helsinki. Imaging for representative images was performed with Pannoramic 250 Flash II high throughput bright field scanner (3DHISTECH) with 20X (NA 0.8) objective and Pannoramic Viewer software.

For immunofluorescence stainings 6-8 μm thick sections were deparaffinized and rehydrated, and antigen retrieval was performed with 1mM EDTA 5-8 minutes in the microvawe oven followed by 20 minutes incubation at RT. Thereafter non-specific binding sites were blocked in immunofluoresence (IF) buffer (7.7 mM NaN3, 0.1% bovine serum albumin [BioWest], 0.2% Triton X-100 and 0.05% Tween-20 [Sigma] in PBS) supplemented with 10% normal goat serum (NGS, Gibco) for 30 minutes at room temperature. Next, samples were incubated with primary antibodies (pan-Laminin 1:500, Abcam; K14 1:500, Covance; K8 1:1000, TROMA-1 Developmental Studies Hybridoma Bank; E-cadherin 1:500, BD; pSTAT5a 1:500, Cell Signaling) overnight at +4°C. The next day samples were washed with 3 times with PBS, 5 minutes each, and incubated with secondary antibodies in 10% NGS in IF Buffer for 1-2 hours. Thereafter samples were washed 3 times with PBS, 5 minutes each followed by counterstaining of nuclei with Hoechst33342 (Sigma). Slides were mounted using Immu-Mount (Thermo Scientific) mounting reagent. Images were acquired using Leica SP8 confocal microscope and HC PL APO CS 63x/1.20W objective and LAS AF software.

### Analysis of p-STAT5a positivity

For quantitation of the number of p-STAT5a positive cells, images from 2-3 areas from individual PP2 mammary gland were acquired. Images were converted to greyscale in Fiji, and nuclei from p-STAT5a and hoechst channels were segmented using edge detection by filtering with difference of gaussians method and thresholding in GNU Image Manipulation Program. Particles from tresholded images were counted using Fiji. Average percentage of p-STAT5a positive nuclei was recorded per mammary gland and mean percentage was calculated per genotype.

### FACS analysis and sorting of primary MMECs

Single cell suspension of isolated primary MMECs was resuspended in 0,2 % BSA in Dulbecco’s PBS and the cells were incubated with following primary antibodies for analysis: CD24-APC, CD31-BV421, CD45-BV421, Ter119-BV421 (Becton Dickinson), CD29-FITC (Sigma), Sca1-BV711 and CD49b-PE (BioLegend) or for sorting: CD31-PE, CD45-PE, Ter119-PE, CD24-Pacific Blue (from Becton Dickinson) and CD29-FITC (Miltenyi Biotec). All antibodies were used 1:500 at RT for 30 minutes. Cells were washed with PBS and re-suspended in 0,2 % BSA in Dulbecco’s PBS with Sytox Blue or 7-AAD (LifeTechnologies, Thermo Fisher) to exclude dead cells. Sorting was performed with FACSAria Fusion (Beckton Dickinson). FlowJo V10 was used for post-analysis of sorted cells.

### RNA isolation and quantitative PCR

RNA isolation was performed using RNeasy isolation kit (Qiagen; from cell cultures) combined with On-Column DNase digestion (Qiagen) or using Trizol (LifeTechnologies, Thermo Fisher; from FACS sorted cells). Trizol isolated RNA was treated with DNase I (Thermo Fisher) according to manufacturer’s protocol. cDNA synthesis was performed with Revert Aid cDNA synthesis kit (Thermo Fisher) starting with 500 ng of RNA. Quantitative PCR reaction was performed with Power SYBR green master mix (Applied Biosystems) and BioRad CFX384 Touch Real-Time PCR detection system. Data was analyzed using BioRad CFX Manager program. Relative mRNA amounts were assayed by comparing PCR cycles to GAPDH using the ddCT method and normalizing to control samples. The following primers were used:

hGAPDH fwd: AAGGTCGGAGTCAACGGAT

hGAPDH rev: TTGATGACAAGCTTCCCGTT

hKRT8 fwd: CAGAAGTCCTACAAGGTGTCCA

hKRT8 rev: CTCTGGTTGACCGTAACTGCG

(Harvard Primer Bank ID: 372466576c1)

hSEMA3B fwd: ACATTGGTACTGAGTGCATGAAC

hSEMA3B rev: GCCATCCTCTATCCTTCCTGG

(Harvard Primer Bank ID: 54607087c1)

mGAPDH fwd: AAGGTCGGAGTCAACGGATT

mGAPDH rev: TTGATGACAAGCTTCCCGTT

mKrt8 fwd: TCCATCAGGGTGACTCAGAAA

mKrt8 rev: CCAGCTTCAAGGGGCTCAA

(Harvard Primer Bank ID: 52789a1)

mKrt14 fwd: AGCGGCAAGAGTGAGATTTCT

mKrt14 rev: CCTCCAGGTTATTCTCCAGGG

(Harvard Primer Bank ID: 21489934c1)

mWap fwd: CGCTCAGAACCTAGAGGAACA

mWap rev: CGGGTCCTACCACAGGAAAC

(Harvard Primer Bank ID: 6755989a1)

mB-cas fwd: GGCACAGGTTGTTCAGGCTT

mB-cas rev: AAGGAAGGGTGCTACTTGCTG

(Harvard Primer Bank ID: 6753538a1)

mCsn3 fwd: AACTGCCGTGGTGAGAAGAAT

mCsn3 rev: AAAGATGGCCTGTAGTGGTAGTA

(Harvard Primer Bank ID: 145966881c1)

mWnt4 fwd: GTACCTGGCCAAGCTGTCAT

mWnt4 rev: CTTGTCACTGCAAAGGCCAC

mLama5 fwd: TTGGTGCGTGTGGAGCGGGC

mLama5 rev: ACTAGGAAGTGCCAGGGGCAG

### Tissue culture

HMEC FL2 cell line was cultured as previously described (Katajisto et al. 2015) in Mammary epithelial basal medium (MEGM; Lonza). For Laminin preadhesion assays, glass coverslips or 12-well plates (Nunc) were coated with 50 μg/ml of Laminin-111 or -521 (BioLamina) at + 4°C overnight before seeding cells. For mammosphere culture, cells were cultured in 1% Methylcellullose (Sigma) in MEGM growth media on 96-well ultra-low attachment plates (Corning) for 7 days in indicated densities. Aggregates showing >2 cells were considered as spheroids and counted. 293fT cells (Thermo Fisher) were grown in DMEM (Sigma) supplemented with 10 % FCS (Gibco), Penicillin/Streptomycin (Orion/Sigma) and 2 mM glutamine (Sigma).

### Primary cell isolation and culture

Primary mammary epithelial cells (MMECs) were isolated from 10-16 weeks old virgin female mice unless otherwise stated. Mammary glands #3-#5 were dissected, and the lymph node in #4 glands was removed and the glands were finely chopped. Tissue was incubated with 0.01 mg of Collagenase A (Sigma) per 1 g of tissue in DMEM/F12 growth media (Life Technologies) containing 2,5% FCS, 5 μg/ml insulin, 50 μg/ml gentamicin and 2 mM glutamine in gentle shaking (120 rpm in environmental shaker) at 37 °C for 2-2,5 hours. The resulting cell suspension was then first centrifuged 400 RCF for 10 minutes and consecutively pulse centrifuged 3-5 times 400 RCF to get a preparation free of other cells than MMEC organoids. Next organoids were trypsinized with 0.05% Trypsin-EDTA (DIFCO, J.T.BAKER) for 5-10 minutes to obtain smaller organoid units and drained trough 70 μm cell strainer (Beckton Dickinson) and resuspended in MMEC growth media (DMEM/F12 media containing 5 μg/ml insulin, 1 μg/ml hydrocortisone, 10 ng/ml mouse EGF, 2 mM glutamine, 50 μg/ml gentamycin and penicillin and streptomycin (all from Sigma) supplemented with 10% FCS (Gibco) for further experiments. For mammosphere culture, cells were cultured in 1% Methylcellullose (Sigma) in DMEM/F12 growth media on 96-well ultra low attachment plates (Corning) for 14 days in indicated densities. Aggregates showing >2 cells were considered as spheroids and counted.

### RNA sequencing and data analysis

RNA was isolated using RNeasy isolation kit (Qiagen) as described above. Total RNA was subjected to quality control with Agilent Tapestation according to the manufacturer’s instructions. To construct libraries suitable for Illumina sequencing the Illumina TruSeq Stranded mRNA Sample preparation protocol which includes mRNA extraction, cDNA synthesis, ligation of adapters and amplification of indexed libraries was used. The yield and quality of the amplified libraries were analysed using Qubit by Thermo Fisher and the Agilent TapeStation. The indexed cDNA libraries were next normalised and combined and the pools were sequenced on the Illumina HiSeq 2000 for a 50-cycle v3 sequencing run generating 50 bp single-end reads. Basecalling and demultiplexing was performed using CASAVA software with default settings generating Fastq files. The resulting Fastq files were passed to STAR for alignment to the human reference genome (hg38) and read counting of annotated genes. The reference genome and annotations were obtained from UCSC. The gene counts were then imported to R/Bioconductor. Reads were TMM normalized and analysis of differential gene expression was carried out with generalized linear models in EdgeR (McCarthy et al., 2012). For the analysis, only genes that had at least one count per million in four or more samples were considered. Gene set enrichment analysis (GSEA) was carried out with default parameters using log2 fold change pre-ranked gene lists (all genes with >1 CPM) of LM-521 grown cells vs cells grown on uncoated dishes. Tested gene sets included Hallmark gene set collection and gene sets upregulated and downregulated in basal HMECs (M5505 and M13422; Huper and Marks 2007) which are available from Molecular Signatures DataBase (MSigDB http://www.broad.mit.edu/gsea/.msigdb/msigdb_index.html).

### Fat pad reconstitution assay

The fat pad reconstitution assay was performed as described previously (Welm et al. 2008). MMECs for reconstitution were isolated from 10-16 week-old wild type C57BL6/RccHsd donor mice as described above. 3-week-old female C57BL6/RccHsd mice were anesthetized using 2-2,5% Isoflurane (Baxter) and the anterior part of #4 mammary gland, lymph node and bridge to #5 gland were surgically removed. To the remaining fat pad were injected indicated amount of MMECs (1*10^3-10^5) in 10 μl volume using Hamilton syringe. The wound was sutured using wound clips (Autoclip Physicians kit, Becton Dickinson), which were removed 1 week after the operation. The fat pad reconstituted mice were sacrificed 8 weeks after reconstitution.

### Lentiviral virus production and infection

Lentiviruses were produced in 293fT cells (Thermo Fisher) grown in DMEM (Sigma) supplemented with 10 % FCS (Gibco), Penicillin/Streptomycin (Orion/Sigma) and 2 mM glutamine (Sigma). Transfections of transfer vector (pLentiCRISPRv2 SCRA, pLentiCRISPRv2 LAMA5) and packaging plasmids (CMV-VSVg and Delta8.9) were performed using Lipofectamine 2000 (Invitrogen) according to manufacturer’s instructions. To concentrate lentiviral particles, media from transfected cells was harvested 72 hours post-transfection and concentrated by ultracentrifugation 22 000 rpm for 120 minutes (Sorvall Discovery 90SE Ultracentrifuge). The viral pellet was then resuspended into PBS to achieve approximately 300-fold concentration and let dissolve at + 4 °C for 18 hours. Viral titer was determined using p24 ELISA test measuring viral capsid protein p24 (Perkin Elmer, performed in Biomedicum Functional Genomics Unit). Primary MMECs were infected on 24-well low adhesion plates overnight using multiplicity of infection (MOI) 5 in 800 μl volume and washed with growth media the next day.

### Plasmid construction for CRISPR guides

CRISPR guides were designed to target LAMA5 using the CRISPR Design tool (http://crispr.mit.edu). Following target sequences were used:

LAMA5: CCATCGATGGCACGGAGCGC SCRA: CTAAAACTGCGGATACAATC Oligos with target sequences were designed according to published instructions (Shalem et al. 2014), annealed and cloned into the pLentiCRISPRv2 vector.

### 3D organotypic cell culture

3D organotypic culture was performed in growth factor reduced basement membrane from Engelbreth-Holm-Swarm (EHS) mouse sarcoma (Matrigel^tm^, Becton Dickinson), which was prepared according to manufacturer’s instructions.

Isolated primary MMECs grown on low adhesion plates were trypsinized with 0.05 % Trypsin-EDTA for 5-10 minutes, centrifuged and suspended with liquid Matrigel and plated onto 8-chamberslides approximately 1500 cells/well. Organoids were grown in DMEM/F12 supplemented with ITS media supplement (Sigma), penicillin and streptomycin and 2,5 nM FGF-2 (Sigma). Wnt3a, Wnt4, Wnt5a (all from R&D Biosystems) were used in 100 ng/ml, Rspo1 (R&D Biosystems) in 500 ng/ml and Porcupine inhibitor IWP-2 (Sigma) in 2 μM were added on the starting day of the cultures. Media was refreshed every 3-4 days.

### Immunofluorescence staining and imaging

2D coverslip grown samples were fixed with 4% PFA for 10 minutes at room temperature. After fixation, cells were washed with PBS and permeabilized with 0.1% Triton-X (Sigma) in PBS for 5 minutes. Next, samples were washed twice with PBS and non-specific binding sites were blocked with 10% FCS in PBS for 30 minutes, and thereafter incubated for 1 hour at RT with primary antibodies (*α*-tubulin, Cell Signaling 1:500, Keratin 8, Developmental Studies Hybridoma Bank 1:300; Keratin 14, Covance 1:300) diluted in blocking solution. Following incubation, samples were washed three times with PBS and then incubated for 30 minutes at RT with appropriate Alexa Fluor (Life Technologies) secondary antibodies (either Alexa 488 or Alexa 594). Finally, samples were washed for 3 times with PBS followed by counterstaining of nuclei with Hoechst33342 (Sigma).

3D organoids were fixed with 2% PFA for 20 minutes at room temperature and thereafter washed with PBS. Epithelial structures were permeabilized with 0.25% Triton X-100 in PBS for 10 minutes at + 4°C, and thereafter washed with PBS. The non-specific binding sites were blocked with 10% normal goat serum (Gibco) for 1-2 hours. The primary antibodies (Keratin 14, Covance 1:300; Ki-67, Abcam 1:500) was incubated in the blocking solution overnight at +4°C. Following the incubation, structures were washed three times with IF buffer, 15 minutes each wash, and then incubated with appropriate Alexa Fluor secondary antibody diluted in blocking solution. After 40-50 minutes incubation at RT, the structures were washed with IF buffer as before and the nuclei were counterstained with Hoechst33342.

All slides were mounted using Immu-Mount (Thermo Scientific) mounting reagent. Images were acquired using Leica TCS SP5 confocal microscope and HCX PL APO CS 63x glyserol (NA 1.3) objective or Leica TCS SP8 STED 3X CW 3D confocal with HC PL APO 63x water (NA 1.20) motCORR CS2 objective and LAS AF software.

### Electron microscopy

Cells were processed for scanning electron microscopy (SEM) in the Electron Microscopy Unit of Institute of Biotechnology at University of Helsinki. Cells were fixed with 2 % glutaraldehyde in 100 mM Na-Cacodylate buffer (pH 7.4) for 1-2 h at room temperature and washed subsequently twice with 0.1M NaCac buffer 5 minutes each. Thereafter, samples were osmicated with 1 % OSO4 in 0.1M NaCac buffer for 1 h and washed with twice with 0.1M NaCac 5 minutes each wash, and then three times with dH2O 10 minutes each wash Samples were then dehydrated and dried using using methylhexadisilatzane (Fluka) overnight prior platinum coating. SEM images were acquired using FEI Quanta 250 Field Emission Gun SEM.

### Quantitation of cell proliferation and growth

For quantitation of cell proliferation cells were seeded on Laminin coated coverslips and grown for 48 hours. For quantification of cell proliferation, cells were treated with 10 μM 5-ethynyl-2’-deoxyuridine (EdU) for 2h and thereafter fixed with 4 % PFA. Immunostaining to detect EdU positive cells was perfomed using Click-IT EdU Alexa Fluor 647 Imaging Kit (ThermoFisher) according to manufacturer’s instructions. Before mounting, nuclei were counterstained with Hoechst33258 (Sigma). For quantification of cell growth cells were seeded on Laminin coated multiwell plates and placed in Cell-IQ (Chip-Man Tecnologies) cell culture platform. Cells from multiple locations within wells were imaged with phase contrast every 24 hours. Cell number per timepoint was recorded from the images using Fiji software.

### Quantification and statistical analysis

Data is presented as mean ± SD from at least three independent experiments or three individual animals quantitated, unless otherwise stated in the figure legend. Unpaired, two-tailed Student’s t tests was used to compare two groups. Statistic tests were performed using GraphPad Prims 6.

## Data availability

RNAseq data will be deposited to GEO and fully available pending acceptance of the manuscript for publication.

## ACKNOWLEDGEMENTS

J. Bärlund, M. Simula, E. Tiilikainen and T. Raatikainen are thanked for excellent technical assistance. H. Hamidi is thanked for language editing of the manuscript. We thank all the members of Katajisto laboratory for comments and discussion. Light Microscopy Unit (LMU) at the Institute of Biotechnology, University of Helsinki and Live Cell Imaging Unit/Nikon Center of Excellence at Department of Biosciences and Nutrition, Karolinska Institutet, are acknowledged for assistance with the microscopy. The study was funded by grants from: European Research council (ERC, #677809 P.K. and #615258 J.I.), Academy of Finland (#266869 P.K., #304591 P.K., #312517 J.I.), Knut and Alice Wallenberg Foundation (KAW 2014.0207), Center for Innovative Medicine (CIMED), Cancerfonden, Sigrid Juselius Foundation, and Finnish Cancer Society.

J.I.E. designed and performed experiments, analyzed the data and wrote the paper; H.C., L.B., A.R, N.P., J.D., and P.M. performed experiments, M.P. provided new reagents, J.K., and J.I. provided new reagents and gave conceptual advice, P.K. designed experiments, analyzed the data and wrote the paper.

The authors declare that they have no conflict of interest.

