## Supplementary figures and images for "Laminin alpha 5 is Necessary for Mammary Epithelial Growth and Function by Maintaining Luminal Epithelial Cell Identity"

### Supplemental Figure 1

Figure S1. Englund JI et al. (related to Figure 1)

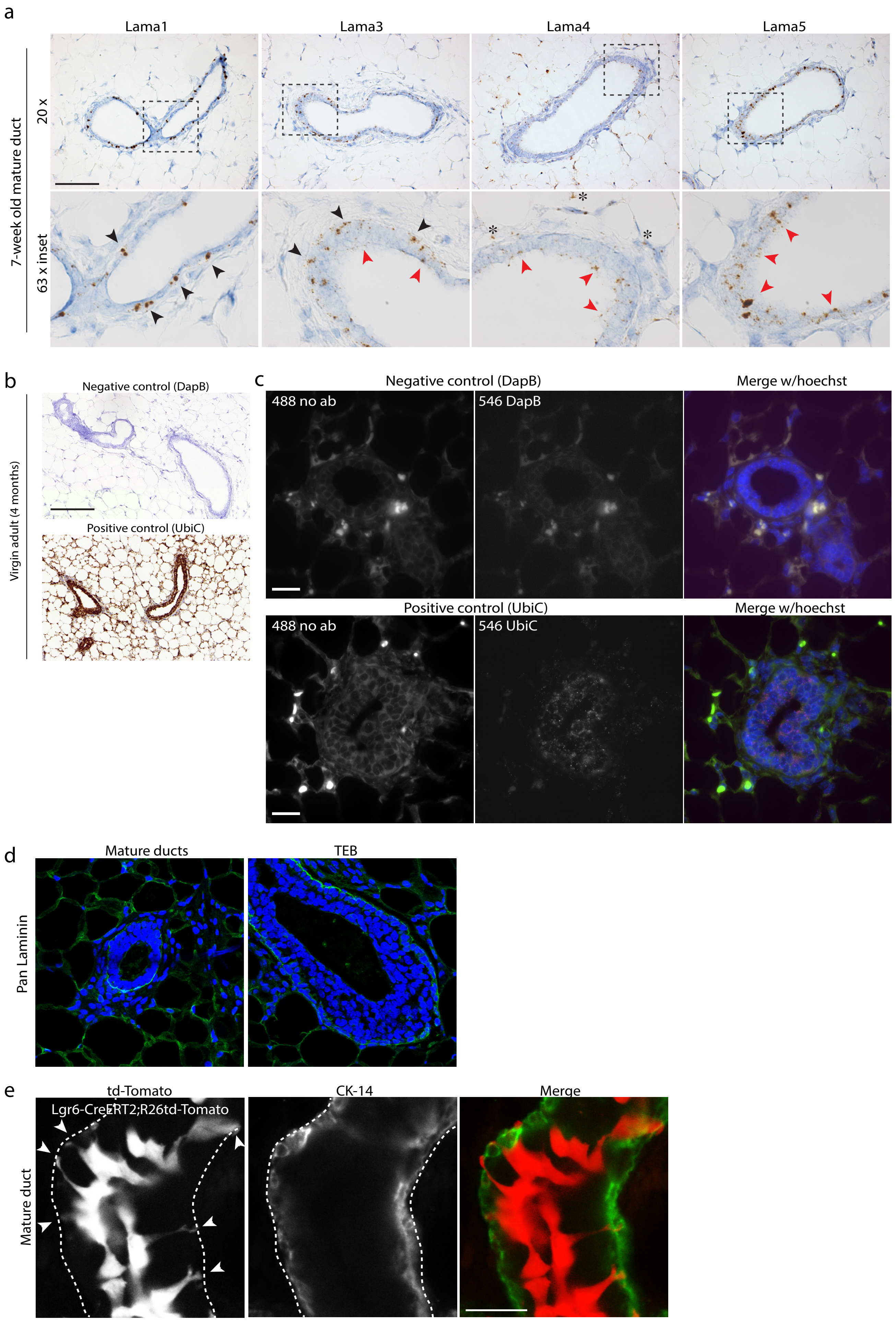

### Supplemental Figure 2

Figure S2. Englund JI et al. (related to Figure 2)

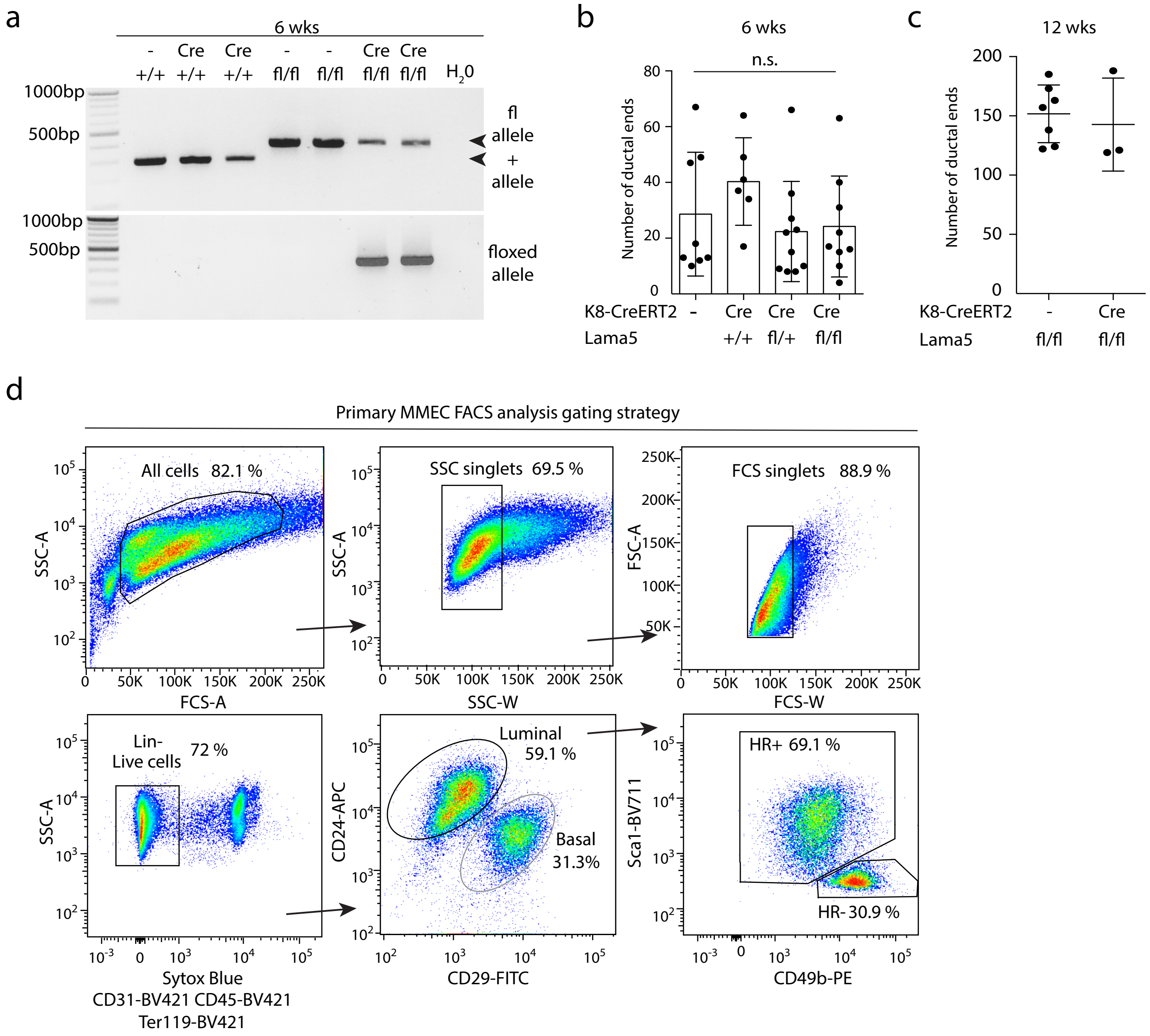

### Supplemental Figure 3

Figure S3. Englund JI et al. (related to Figure 4)

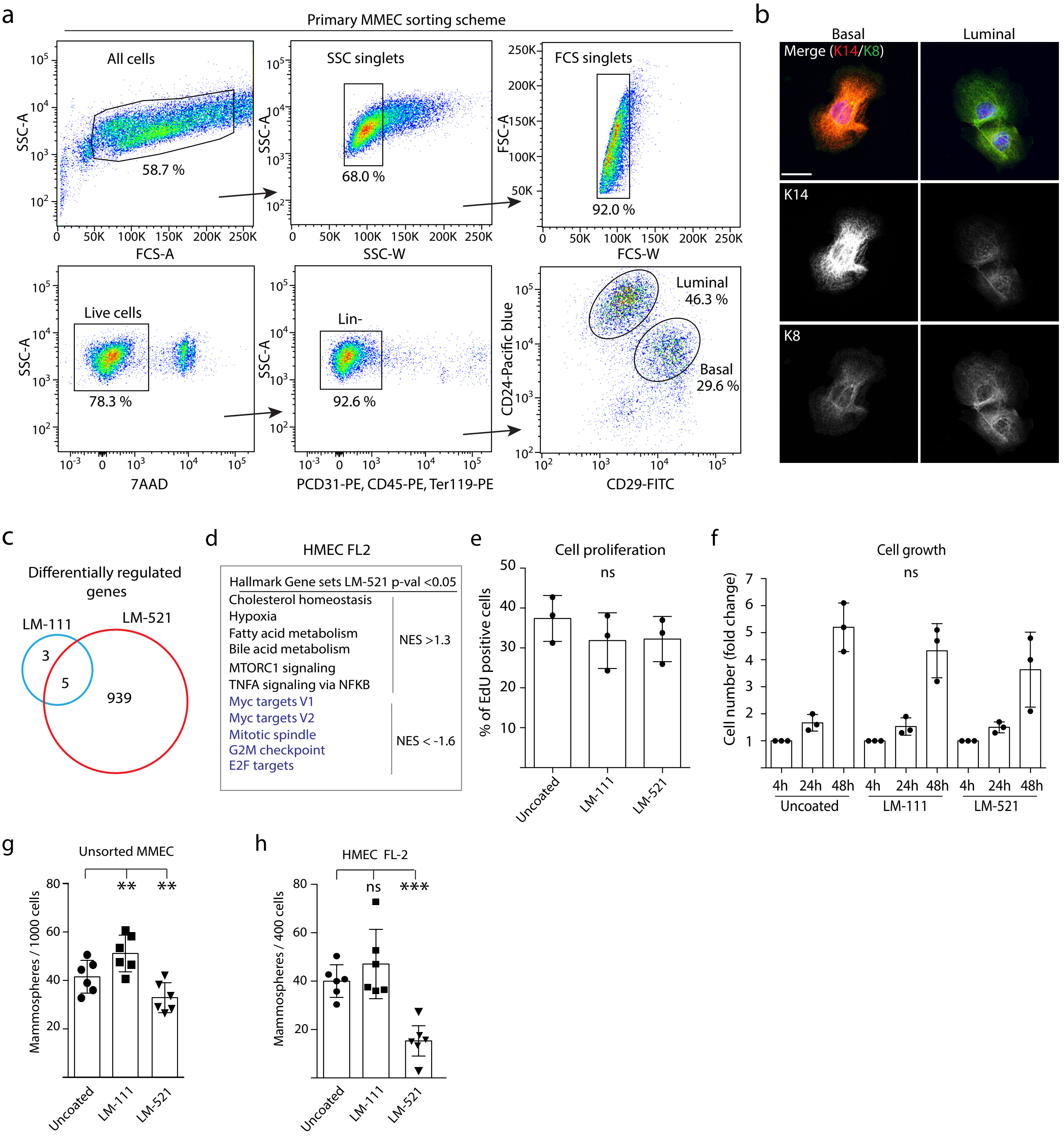

### Supplemental Figure 4

Figure S4. Englund JI et al. (related to Figure 5)

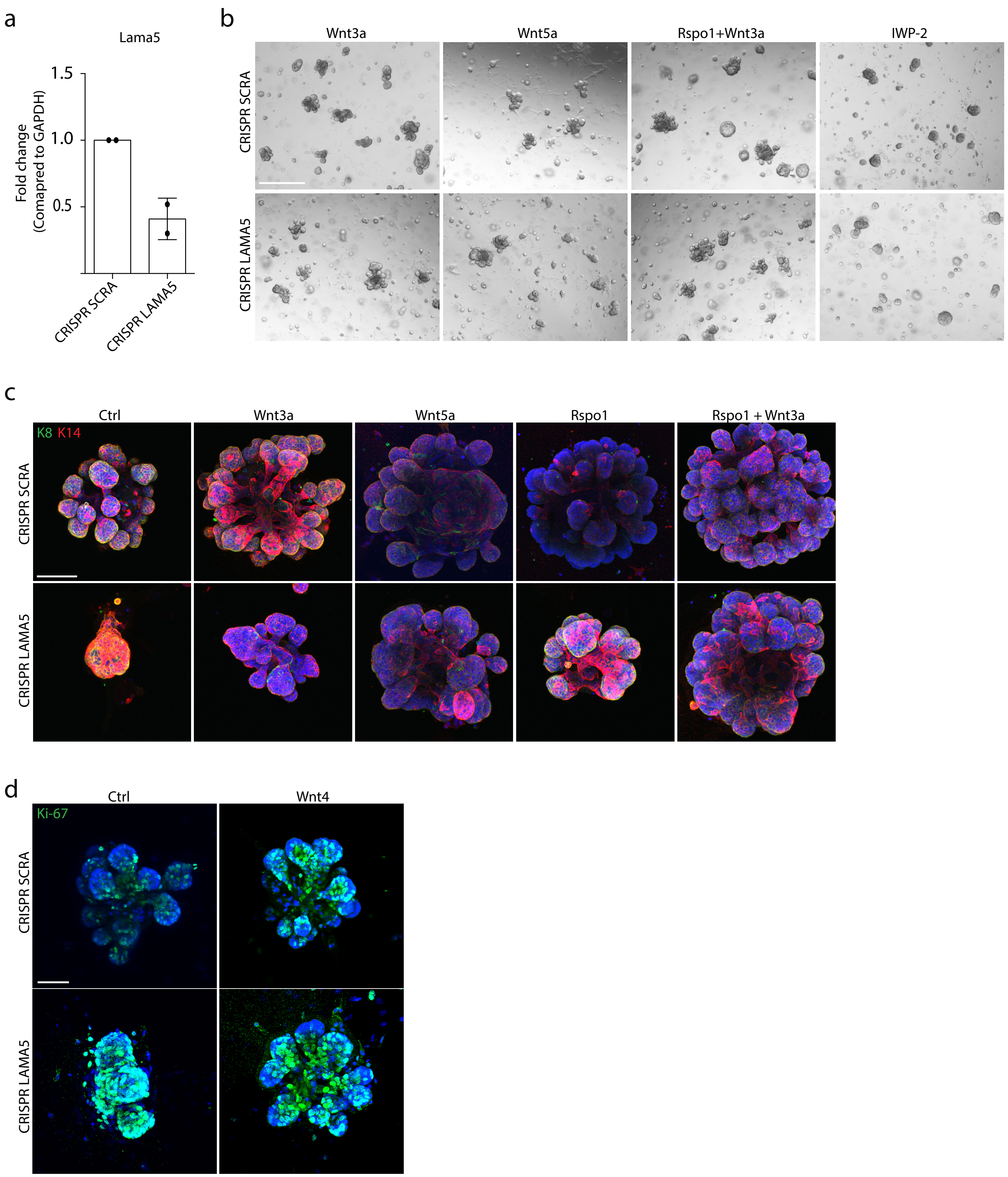
