## Supplemental Figure legends for "Laminin alpha 5 is Necessary for Mammary Epithelial Growth and Function by Maintaining Luminal Epithelial Cell Identity"

**Figure S1. (Related to Figure 1) Laminin $\alpha$ chain expression in the mammary gland.** a) RNA *in situ* hybridization performed with probes for *Lama1*, *3*, *4* and *5* on mature ducts of 7-week old pubertal mammary glands. Expression by basal or luminal cells is indicated by black and red arrowheads, respectively. Black asterisks mark expression in stromal cells. Scale bars 50 $\mu$m, 20 $\mu$m (insets). b) Representative images of RNA in situ hybridization negative control (DapB) and positive control (UbiC) for chromogenic staining in young adult (4 months old) mammary gland. Scale bar 50 $\mu$m. c) Representative images of RNA in situ hybridization negative control (DapB) and positive control (UbiC) for immunoluorescence in 7-week old mammary gland. Scale bar 20 $\mu$m d) Immunofluorescence images of 7-week old mature mammary ducts and TEBs with Pan-Laminin antibody. Scale bar 50 $\mu$m. e) A representative image of luminal cells (red) in tamoxifen-treated 4-week old *Lgr6-CreERT2*;*Rosa26-tdTomato* mammary epithelium immunostained to visualize Keratin 14 (green). Dotted line marks the interface between BM and the epithelium. Note the extensions by bright tdTomato positive luminal cells to the BM (white arrowheads). Scale bars 20 $\mu$m.

**Figure S2. (Related to Figure 2) Laminin $\alpha$5 is required for growth of pubertal mammary epithelium.** a) Genomic PCR of DNA isolated from mammary glands of 6-week old mice treated with tamoxifen at 3-weeks of age. Upper gel shows PCR products of Lama5$^{+/+}$ (296 bp) or Lama5$^{fl}$ allele (460 bp) and lower gel products of recombined Lama5$^{fl}$ allele (250 bp, predicted). b) Quantification of the number of ductal ends in 6-week old transgenic mice treated as in a. Analyzed from #4 mammary glands, data show mean $\pm$ SD. Two-tailed student's t-test was used to compare the indicated groups. c) Quantification of the number of ductal ends in 12-week old transgenic mice treated with tamoxifen at 8-weeks of age. d) Representative FACS plots and gating strategy for separation of basal and luminal primary MMECs and HR- and HR+ luminal subpopulations. Same CD24/CD29 plot from *Lama5$^{+/+}$;K8-CreERT2* control animal with luminal and basal sorting gates is shown in Figure 2j.

**Figure S3. (Related to Figure 4) Adhesion to specific laminins affects luminal and basal gene expression.** a) Representative FACS plots and gating strategy for the separation and sorting of luminal and basal primary MMECs. b) Representative immunofluorescence images of K8 and K14 stained basal and luminal cells FACS sorted as in a). Scale bar 20 μm. c) Venn diagram showing differentially expressed genes in HMECs grown on indicated laminins and compared to uncoated control. d) Top Hallmarks gene sets in Gene Set Enrichment Analysis (GSEA; MSigDB collection) showing positive (black) or negative (blue) correlation with gene expression signature in LM-521 grown cells with nominal p-value <0.05. e) Quantitation of cell proliferation using EdU. Percentages indicate average number of EdU-positive cells on the indicated laminins from three independent experiments. Data show mean ± SD. Two-tailed student's t-test was used to compare results to uncoated control. f) Quantitation of cell growth from phase contrast live cell imaging of cells on indicated laminins normalized to 4 hour timepoint. Data show mean ± SD. Two-tailed student's t-test was used to compare results to uncoated control. g) Mammosphere forming frequency of unsorted MMECs pre-adhered on the indicated laminins for 48h 7 days prior. Data show mean ± SD from independent experiments. h) Mammosphere forming frequency of basal HMECs pre-adhered on the indicated laminins for 48h 7 days prior. Data show mean ± SD from independent experiments.

**Figure S4. (Related to Figure 5) *Lama5* deleted luminal MECs exhibit defective Wnt signaling.** a) qRT-PCR analysis of Lama5 mRNA expression in MECs infected with lentiviruses carrying either LAMA5 or SCRA CRISPR guides compared to GAPDH. Quantification shows average of technical repeats from one experiment. N=2 biological replicates in both SCRA and LAMA5. b) Primary MMECs carrying either LAMA5 or SCRA CRISPR guides grown in Matrigel in the presence of 2.5 nM FGF-2 and either with Wnt3a, Wnt5a, Rspo1 and Wnt3a or IWP-2 for 7 days. Scale bar 200 μm. c) Representative immunofluorescence images of LAMA5 or SCRA CRISPR carrying MMECs and as in b and stained for K8 and K14. Scale bar 100 μm. d) Representative immunofluorescence images of LAMA5 or SCRA CRISPR carrying MMECs and as in b and stained for Ki-67. Scale bar 50 μm.
